# Staged developmental mapping and X chromosome transcriptional dynamics during mouse spermatogenesis

**DOI:** 10.1101/350868

**Authors:** Christina Ernst, Nils Eling, Celia P Martinez-Jimenez, John C Marioni, Duncan T Odom

**Affiliations:** European Molecular Biology Laboratory, European Bioinformatics Institute, (EMBL-EBI), Wellcome Genome Campus, Hinxton, Cambridge, CB10 1SD, United Kingdom; University of Cambridge, Cancer Research UK Cambridge Institute, Robinson Way, Cambridge, CB2 0RE, United Kingdom; Wellcome Sanger Institute, Welcome Genome Campus, Hinxton, Cambridge, CB10 1SA, United Kingdom; German Cancer Research Center (DKFZ), Division Signaling and Functional Genomics, 69120, Heidelberg, Germany

## Abstract

Understanding male fertility requires an in-depth characterisation of spermatogenesis, the developmental process by which male gametes are generated. Spermatogenesis occurs continuously throughout a male’s reproductive window and involves a complex sequence of developmental steps, both of which make this process difficult to decipher at the molecular level. To overcome this, we transcriptionally profiled single cells from multiple distinct stages during the first wave of spermatogenesis, where the most mature germ cell type is known. This naturally enriches for spermatogonia and somatic cell types present at very low frequencies in adult testes. Our atlas, available as a shiny app (https://marionilab.cruk.cam.ac.uk/SpermatoShiny), allowed us to reconstruct the three main processes of spermatogenesis: spermatogonial differentiation, meiosis, and spermiogenesis. Additionally, we profiled the chromatin changes associated with meiotic silencing of the X chromosome, revealing a set of genes specifically and strongly repressed by H3K9me3 in the spermatocyte stage, but which escape post-meiotic silencing in spermatids.

## INTRODUCTION

Sexual reproduction in eukaryotes drives evolution and adaptation (McDonald et al., 2016). It requires the generation of haploid gametes that upon fusion combine their genetic material and develop into a diploid organism. Gametogenesis is a tightly regulated developmental process that differs dramatically between males and females to generate sperm and eggs (Spiller et al., 2017).

In spermatogenesis, spermatogonial stem cells undergo a unidirectional differentiation programme to form mature spermatozoa. Spermatogenesis occurs in the epithelium of seminiferous tubules in the testis and is tightly coordinated to ensure the continuous production of mature sperm cells throughout the reproductive lifespan of an animal. In the mouse, this differentiation process initiates with the division of a spermatogonial stem cell (SSC or A_single_) to form first a pair, and then a connected chain, of undifferentiated spermatogonia (A_paired_ and A_aligned_) (Oakberg, 1971; de Rooij, 1973). These cells have competency to undergo spermatogonial differentiation, which involves six transit-amplifying mitotic divisions generating A_1-4_, Intermediate, and B spermatogonia, which then give rise to pre-leptotene spermatocytes (pL) (de Rooij and Russell, 2000).

Pre-leptotene spermatocytes then commit to meiosis, a specialised cell division programme that consists of two consecutive cell divisions without an intermediate S phase to produce haploid cells. In contrast to mitosis, meiosis includes programmed DNA double strand break (DSB) formation, homologous recombination, and chromosome synapsis (Marston and Amon, 2004). To accommodate these events, prophase of meiosis I is extremely prolonged, lasting several days in males, and can be divided into four substages: leptonema (L), zygonema (Z), pachynema (P) and diplonema (D). Following the two consecutive cell divisions, haploid cells known as round spermatids (RS) are produced, and then undergo a complex differentiation programme called spermiogenesis to form mature spermatozoa (Oakberg, 1956a).

Spermatogenesis takes place in a highly orchestrated fashion, with tubules periodically cycling through twelve epithelial stages defined by the combination of germ cells present (Oakberg, 1956a). The completion of one cycle takes 8.6 days in the mouse, and the overall differentiation process from spermatogonia to mature spermatozoa requires approximately 35 days (Oakberg, 1956b). Thus, four to five generations of germ cells are present within a tubule at any given time. The continuity of this differentiation process and the gradual transitions between spermatogenic cell types have made the isolation and thus the molecular characterisation of individual sub-stages during spermatogenesis difficult.

To elucidate the molecular genetics of germ cell development, we have used an unbiased droplet-based single-cell RNA-Sequencing (scRNA-Seq) approach to capture the continuum of spermatogenic cell populations in the adult testis. We used these data to characterize the complex and dynamic transcriptional profile of spermatogenesis at high-resolution. To confidently identify and label cell populations throughout the developmental trajectory, we profiled cells from juvenile testes during the first wave of spermatogenesis. In juveniles, spermatogenesis has only progressed to a defined developmental stage, and therefore allowed us to unambiguously identify the most mature cell type by comparison with adult. Furthermore, by profiling juvenile samples in which the cell type composition within tubules is heavily biased towards somatic cells and spermatogonia, we obtain the molecular signatures of poorly-characterized cell populations. For spermatogonia, this allows us to visualise and profile their dynamic differentiation process at high resolution.

Similarly, we dissected the gene expression heterogeneity within spermatocytes and spermatids, the cell types undergoing meiosis and spermiogenesis, respectively. Our analyses revealed a large number of genes with previously described roles in spermatogenesis, but also allowed us to uncover novel genes that are likely to play important roles during germ cell development and thus relevant for male fertility.

Finally, we focused our attention on the inactivation and reactivation of the X chromosome, which is subject to transcriptional silencing as a consequence of asynapsis (Turner, 2007). By combining bulk and single-cell RNA-Seq approaches, we identified genes that show *de novo* activation in post-meiotic spermatids with distinct temporal expression patterns. Our data revealed that *de novo* activated escape genes carry distinct chromatin signatures with high levels of repressive H3K9me3 in spermatocytes. Overall, our study presents an in-depth characterization of mouse spermatogenesis and provides new insights into the epigenetic regulation of X chromosome reactivation in post-meiotic spermatids.

## RESULTS

### High-resolution profiling by single-cell RNA-seq captures the unidirectional differentiation of germ cells during adult spermatogenesis

Spermatogenesis is a recurrent differentiation process that produces male gametes within testicular seminiferous tubules (**Fig. 1A**). The seminiferous epithelium in the mouse is classified into twelve distinct stages, based on the combination of cell types present (**Fig. 1B**). Tubules cycle asynchronously and continuously through these stages and adult testis contain tubules in every possible epithelial stage (**Fig. 1A** **and B; Fig. S1A**).

Using multiple functional genomics approaches, we characterized the transcriptional programme underlying mouse spermatogenesis with cells isolated from specifically staged juvenile (between postnatal days 6 and 35) and adult (8-9 weeks) C57BL/6J (B6) mice. For all samples, we generated unbiased droplet-based single-cell RNA sequencing (scRNA-Seq) data from whole testis with matched histology. Additionally, for juvenile samples, we generated whole-tissue bulk RNA sequencing, as well as mapping chromatin state in purified cell populations using CUT&RUN (Cleavage Under Targets & Release Using Nuclease) (**Fig. 1C**; **Methods**) (Skene et al., 2018). After quality control and filtering, we retained a total of 42,796 single cells, 30 bulk RNA-Seq libraries and 8 CUT&RUN libraries (**Methods; Table S1**).

After clustering all single cell transcriptomes using an unbiased graph-based approach (**Methods**), we first focused on cells isolated from adult B6 testis to generate a comprehensive map of cell types across spermatogenesis. Using computationally-defined cluster-specific marker genes (**Methods; Table S2**), we identified the following cell types: spermatogonia (based on *Dmrt1* expression, Matson et al., 2010), spermatocytes (*Piwil1*, Deng and Lin, 2002), round and elongating spermatids (*Tex21* and *Tnp1*, respectively, Fujii et al., 2002), as well as the main somatic cell types of the testis, Sertoli (*Cldn11*, Mazaud-Guittot et al., 2010) and Leydig cells (*Fabp3*, Oresti et al., 2013) (**Fig. 1D**). Using a dimensionality reduction technique for visualization (t-distributed Stochastic Neighbour Embedding; **Fig. 1E**), the germ cell types from spermatocytes to elongating spermatids formed a continuum, which recapitulated the known developmental trajectory.

**Figure 1:**
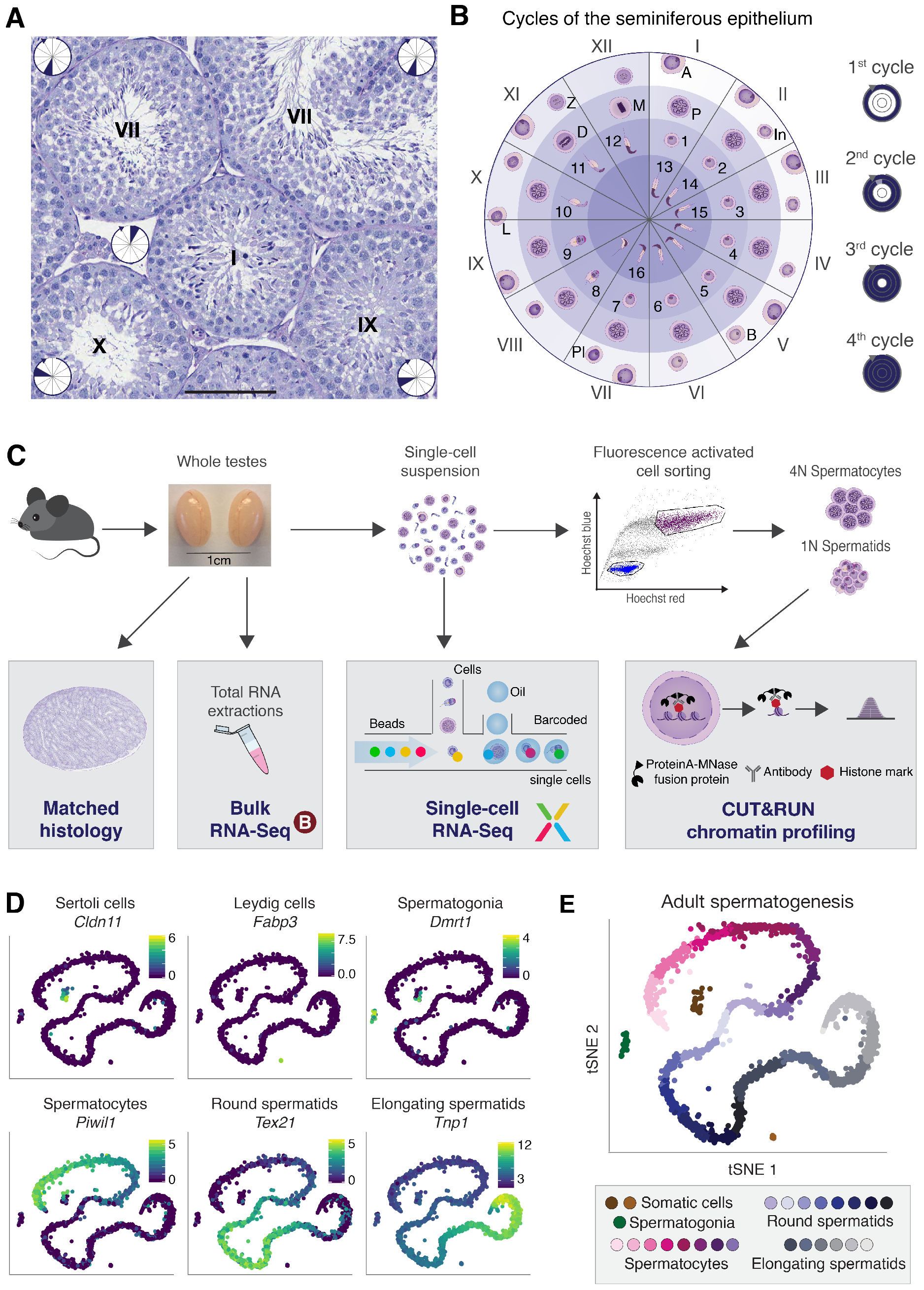
Single-cell RNA-Seq captures a continuum of germ cell types during adult spermatogenesis. **(A)** Periodic Acid Schiff (PAS)-stained testis cross-section showing a number of seminiferous tubules at different epithelial stages (displayed as Roman numerals). Within each tubule, the inset circle refers to the corresponding section in (B). Scale bar represents 100 *μ*m; original magnification 200X. **(B)** Schematic representation of the 12 stages of the seminiferous epithelium in mice. The colour gradient within the circle indicates the differentiation path of germ cells with the layers corresponding to individual cycles of the epithelium. The circle is divided into 12 section, each corresponding to one epithelial stage displaying the characteristic germ cells. Within each section, cells are positioned across the different layers according to their emergence during consecutive cycles, each being 8.6 days apart with more mature cells moving towards the centre. Cell types are labelled as: A - type A spermatogonia (SG), In - intermediate SG, B - type B SG, Pl - preleptotene spermatocytes (SCs), L - leptotene SCs, Z - zygotene SCs, P - pachytene SCs, D - diplotene SCs, M - metaphase I and II, 1-8 round spermatids, 9-16 elongating spermatids. **(C)** Overview of the experimental design yielding bulk RNA-Seq, droplet-based scRNA-Seq and chromatin profiling on FACS-purified cells using CUT&RUN from one testis while using the contralateral testis for matched histology. **(D)** t-distributed stochastic neighbour embedding (tSNE) representation of scRNA-Seq data from adult B6 mice with the colour gradient representing the expression of known marker genes for two somatic cell types and the main germ cell types. The x-and y-axis represent the first and second dimension of tSNE respectively. The colour legend shows log_2_-transformed, normalized expression counts. **(E)** Graph-based clustering (see Methods) identifies different sub-stages within major germ cell populations.

### Staged developmental mapping of the first wave of spermatogenesis defines germ cell identity

Historically, sub-staging of the major cell types within the testis was based on changes in nuclear or cellular morphology (**Fig. S1B**) (Oakberg, 1956a, 1956b). Previous attempts to complement morphology with molecular signatures have been limited to FACS-based and sedimentation assays, where the resolution was unable to differentiate between sub-cell-types (Bastos et al., 2005; Gaysinskaya et al., 2014; Getun et al., 2011; Lam et al., 1970; Meistrich, 1977; Romrell et al., 1976; Soumillon et al., 2013). While a mixture of all spermatogenic cell types co-exists in the adult, the first wave of spermatogenesis during juvenile development is more synchronised. Starting around mouse postnatal day P4, spermatogonia begin to differentiate, forming the first generation of spermatocytes as early as P10, round spermatids by P20, and completing the first wave of spermatogenesis with the production of mature spermatozoa between P30 and P35 (**Fig. 2A**) (Bellvé et al., 1977; Janca et al., 1986; Nebel et al., 1961).

We exploited this first spermatogenic wave to generate thousands of high-resolution single-cell transcriptional profiles of morphologically well-defined cell populations. We sampled multiple time points during the first wave to identify the most mature (differentiated) cell types. Based on known sperm developmental transitions, we chose six time points between P10 and P35 from which to generate single-cell RNA-Seq libraries (**Fig. 2A**). For each juvenile experiment, we found that the population of developing germ cells was strongly enriched at the expected developmental stage, as quantified by the percentage of cells in each cell-type cluster (**Fig. 2B** **and C; Methods; Fig. S2A**). By associating these cell types with the corresponding cell types in the adult trajectory, we unambiguously assigned molecular and histological signatures to cells during adult spermatogenesis.

For instance, at P15 the majority of cells are spermatogonia and spermatocytes progressing through the mid-pachytene stage, in accord with the appearance of the sex body (Turner et al., 2004a). Interestingly, less mature cell types are also present at this time point (and later time points), supporting recent reports that the first wave of spermatogenesis is less synchronized than previously anticipated (Snyder et al., 2010). At P20, we detect an enrichment for spermatocytes, cells undergoing meiotic cell division, and a small group of early round spermatids. This population structure is in line with matched histology, which shows a large number of tubules in late stages IX-XII (**Fig. S2B**) and the first occurrence of early round spermatids (Bellvé et al., 1977).

Spermatids first reach the elongating state between P24 and P26 (Janca et al., 1986). At P25, we observed cells matched to our first ten clusters of spermatids, which we then labelled according to morphologically-defined spermatid substages S1 - S10 (**Fig. 2C**; **Fig. S2A**). At P30 and P35, we observed a relatively even distribution of cells across all groups, closely resembling the adult distribution up to S14, indicating that the first wave of spermatogenesis is complete.

To further validate the identity of the cell clusters, we used bulk RNA-Seq from testis collected during the first wave of spermatogenesis every two days between P6 and P34 (**Fig. 2A**). We matched the bulk and single-cell RNA-Seq data by using a probabilistic classification of the bulk samples based on the cluster-specific marker genes in the adult data (**Methods**; **Fig. 2D**). This confirmed that between P6 - P14 spermatogonia and somatic cells show the highest contribution to the transcriptomic profile. Between P16 and P20 we observed the emergence of spermatocyte-specific gene expression signatures, after which spermatids become the transcriptionally dominant cell type. By P26, spermatids reach the elongating state where transcription is uniformly shut-down due to the beginning of the histone-to-protamine transition (Steger, 1999); following this, changes in RNA content are mostly due to degradation. Bulk transcriptional profiles can therefore only be classified up to S10, because transcription is largely inactive thereafter.

In sum, by combining transcriptional and histological analyses at specific stages during the first wave of spermatogenesis, we assigned transcriptional profiles to specific, morphologically defined germ cell types.

**Figure 2:**
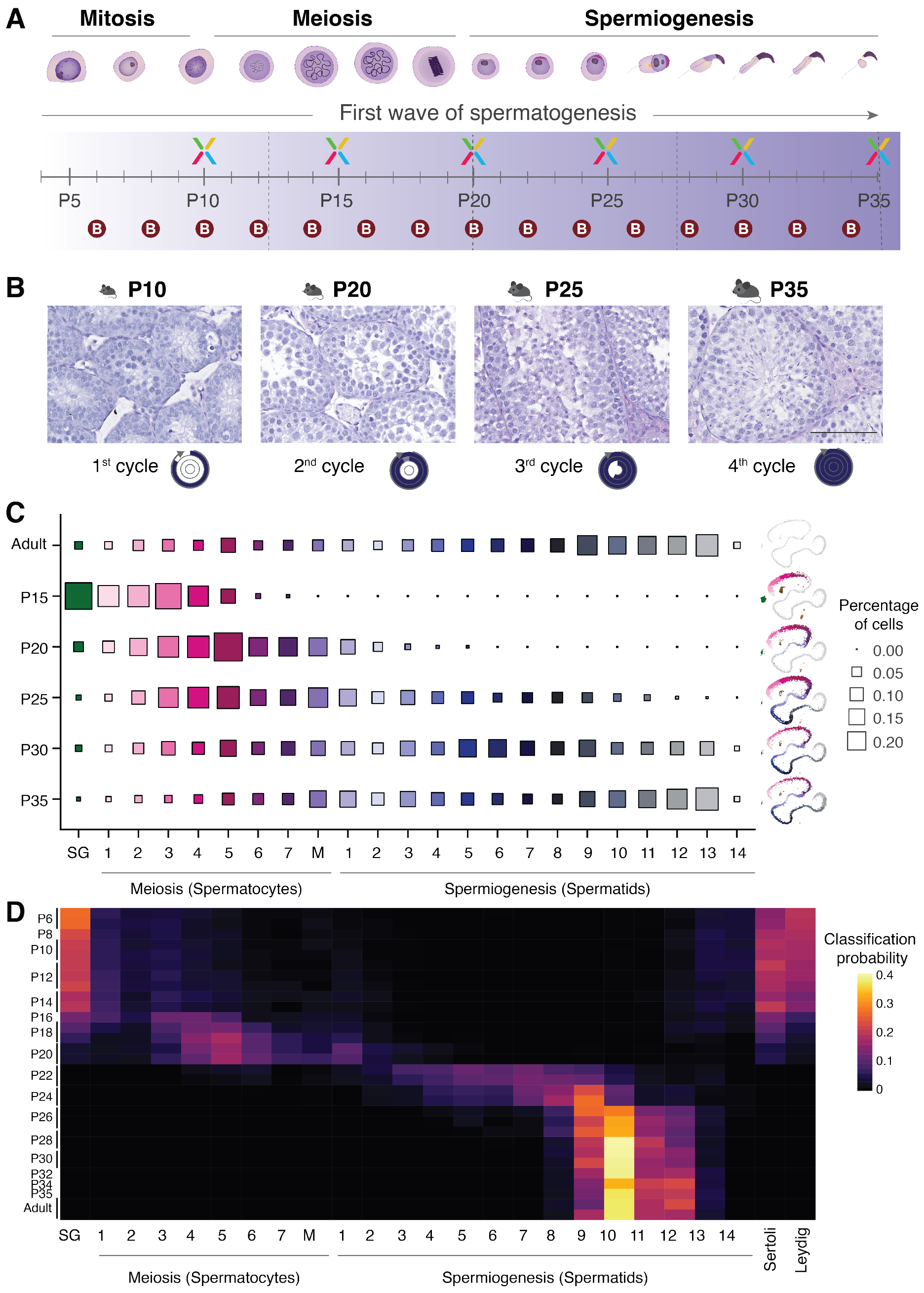
Cell type classification by developmental mapping of time points during the first wave of spermatogenesis. **(A)** Schematic representation of the major germ cell types and their corresponding developmental processes. Spermatogonia differentiate undergoing mitotic cell divisions before forming spermatocytes that divide by meiotic division. Spermatids differentiate throughout spermiogenesis to form mature sperm. The timeline in the lower panel indicates at which point during the first wave of spermatogenesis samples were harvested for the generation of scRNA-Seq (X) or bulk RNA-Seq (B) data. **(B)** Representative images of seminiferous tubules from PAS-stained tissue cross-sections from animals harvested at different postnatal (P) time points during the first wave of spermatogenesis. The approximate timing of the stage and cycle of the tubule is illustrated below in the form of a circle (see Fig. 1B). **(C)** The percentage of cells allocated to each cell cluster for each juvenile and adult sample is represented by the size of squares with the colours corresponding to the clusters depicted in **Fig. 1E**. tSNEs on the right-hand side of each panel (juvenile samples only) illustrate progress through spermatogenesis. SG - spermatogonia, M - metaphase. **(D)** Probabilistic mapping of bulk RNA-Seq libraries to the cell clusters identified in the adult scRNA-Seq data using a random forest approach (see Methods). The colour gradient indicates the probability with which a bulk sample can be assigned to the specific cell cluster.

### Identification of under-represented and transcriptionally silent cell types in adolescent testes

In addition to assigning the dominant cell types in germ cell development, we further exploited the juvenile samples to analyse the highly enriched cell types that are rare in adult testes, including somatic support cells, spermatogonia and the earliest spermatocytes. Through the first wave of spermatogenesis, the ratio between germ cells and somatic cells increases; therefore, to study heterogeneity within the somatic cell population, we focused on the P10 stage, where somatic cells are relatively more frequent. As expected, we readily identified substantial numbers of Sertoli and Leydig cells, which are the main somatic cell types in adult (Griswold, 1998; Haider, 2004). In addition, we newly identified a large population of immature Leydig cells, based on *Dlk1* expression (Lottrup et al., 2014). Furthermore, we detected the cells that form the basal lamina such as peritubular myoid cells (PTM, *Acta2*, Cool et al., 2008), vascular endothelial cells (*Tm4sf1*, Shih et al., 2009), and testicular macrophages (*Cd14*, Kitchens, 2000) (**Fig. S3B**). We then identified specific marker genes for the different somatic cell populations to enable further in-depth molecular characterization of these rare cell types (**Table S3**).

Identifying spermatogonial sub-populations in adult testes is greatly complicated by their rarity relative to other germ cell types (Lukassen et al., 2018). However, during early juvenile development spermatogonia are relatively enriched, which we exploited to further characterize their heterogeneity (**Fig. 3A**). By combining cells from P10 and P15, we obtained 1,186 transcriptional profiles that capture sub-populations during spermatogonial differentiation (**Fig. 3B**). We detected two clusters corresponding to undifferentiated spermatogonia (A_undiff_) based on their expression of *Nanos3* and *Zbtb16* (**Fig. 3B** **and C**) (Buaas et al., 2004; Lolicato et al., 2008). These cells comprise A_s_, A_paired_, and A_aligned_ spermatogonia that decrease in stemness as they divide and gain competency to differentiate (Suzuki et al., 2012). Additionally, these cells express a number of marker genes also detected in undifferentiated human spermatogonial stem cells, such as *Gfra1*, *Bcl6* and *Id4* (**Table S4**) (Guo et al., 2017). Based on the expression of *Stra8* (Stimulated by retinoic acid 8), we can map the point at which spermatogonial differentiation is induced (A_aligned_-to-A_1_ transition), thus marking the beginning of differentiating spermatogonia (A_diff_) (Endo et al., 2015). A_diff_ are marked by the expression of *Sohlh1* (Ballow et al., 2006) and are highly proliferative, generating A_1-4_, Intermediate and B spermatogonia. Late differentiating spermatocytes express *Dmrtb1* (also referred to as *Dmrt6*), which mediates the mitosis-to-meiosis transition and quickly disappears in preleptotene spermatocytes. This latter population shows a second increase in *Stra8* expression levels, which is necessary for initiation of meiosis (**Fig. 3B** **and C**) (Anderson et al., 2008; Endo et al., 2015; Zhang et al., 2014).

The transition between differentiating spermatogonia and spermatocytes is a gradual process that occurs in stage VIII tubules when B spermatogonia divide and form preleptotene spermatocytes (Anderson et al., 2008; Baltus et al., 2006). We were therefore surprised not to observe a continuous differentiation trajectory bridging spermatogonia to spermatocytes (**Fig. 1E**; **Fig. S3C**). One possible explanation is that leptotene and zygotene spermatocytes have decreased transcriptional activity (Kierszenbaum and Tres, 1974; Monesi, 1965), and are thus likely to be classified as empty droplets by the 10X CellRanger pipeline (**Methods**).

To capture these transcriptionally quiescent cells, we applied a newly developed computational method that distinguishes between droplets capturing genuine cells containing small amounts of mRNA **versus** empty droplets containing only ambient mRNA (**Methods**; (Lun et al., 2018)). Applying this approach increased the number of early spermatocytes in all samples and, in particular, identified a population of cells connecting spermatogonia and spermatocytes at the predicted position in the cell trajectory (**Fig. S4A-C**). As expected for leptotene and zygotene spermatocytes, these cells show high mRNA levels for genes involved in synaptonemal complex formation, chromosome synapsis and DNA double-strand break (DSB) formation such as *Sycp1, H2afx* and *Hormad1* (**Fig. S4D; Table S5**) (Daniel et al., 2011; Mahadevaiah et al., 2001; Vries et al., 2005).

In addition to early spermatocytes, droplets with lower transcriptional complexity also captured late condensing spermatids; as mentioned above, these late stages of spermiogenesis are characterized by continuous degradation of RNA after transcriptional shutdown at the round-to-elongating transition (Steger, 1999) (**Fig. S4E**).

By profiling carefully chosen developmental time points and applying new methods to incorporate single cells with lower transcriptional complexity, our results elucidate a more complete view of testicular cell type composition that includes previously under-represented cell types.

**Figure 3:**
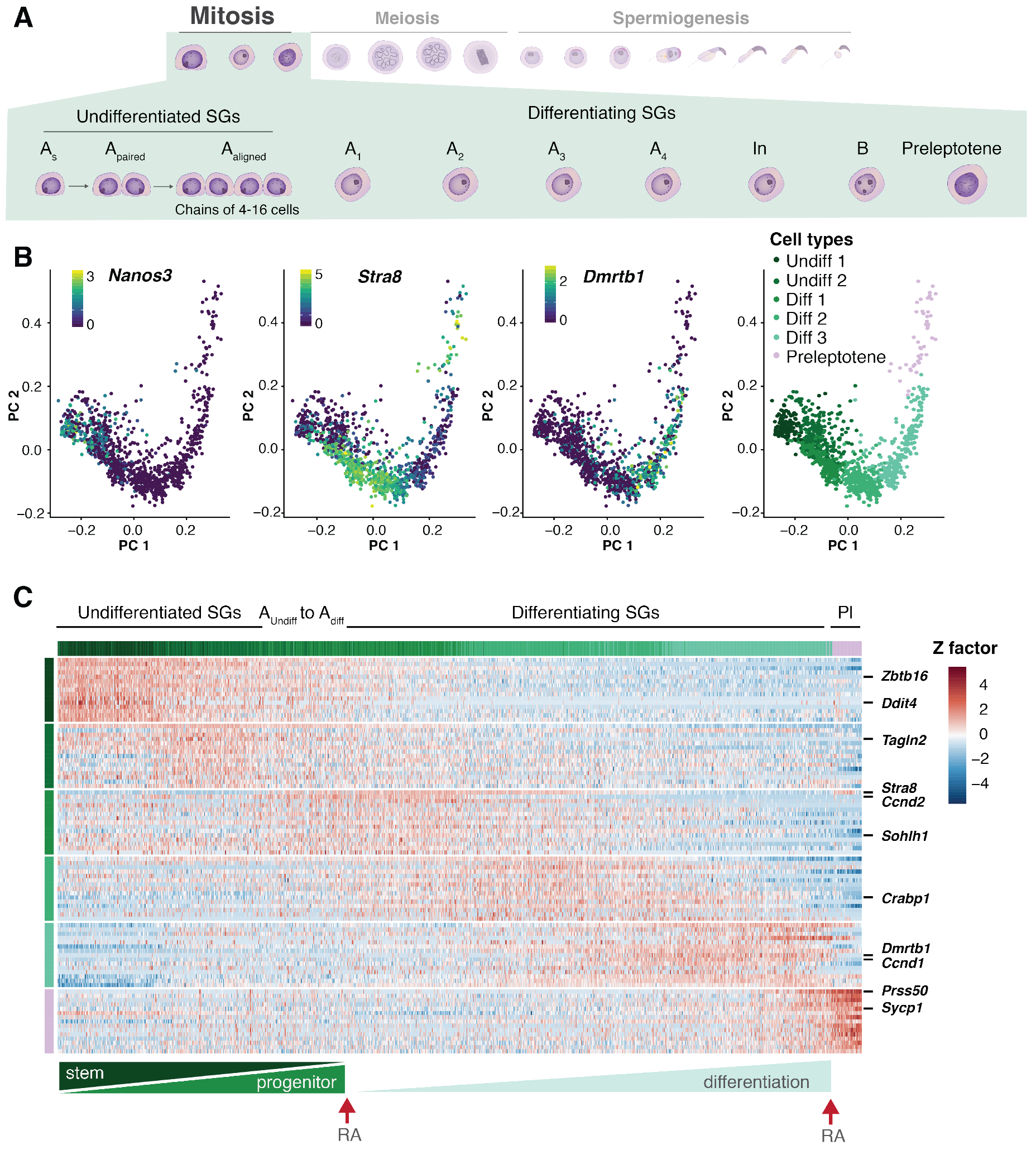
Cellular heterogeneity during spermatogonial differentiation. **(A)** Schematic representation of spermatogonial differentiation including sub-stages of undifferentiated (A_s_, A_paired_, A_aligned_) and differentiating spermatogonia (A_1_, A_2_, A_3_, A_4_, In, B) (SGs) as well as Preleptotene (Pl). **(B)** Sub-structure detection in spermatogonia isolated from P10 and P15 animals. PCA was computed on batch-corrected transcriptomes (see Methods). The first three panels represent expression of known marker genes for undifferentiated (Undiff, *Nanos3*) and differentiating (Diff, *Stra8* and *Dmrtb1*) spermatogonia. The colour scale shows log_2_-transformed, normalized counts. The last panel overlays cluster identity by sub-clustering batch-corrected transcriptomes of spermatogonia (see Methods). **(C)** Scaled, normalized expression counts of the top 15 marker genes per cell cluster. Column and row labels represent the cell clusters identified in the last panel of (B). The lower bar indicates the gradual differentiation from undifferentiated spermatogonia to preleptotene cells driven by two retinoic acid (RA) signals.

### High-resolution characterization of male meiosis

The mitotic expansion of spermatogonia produces large numbers of spermatocytes, which then undergo male meiosis where two consecutive cell divisions give rise to four haploid spermatids. In contrast to mitotic cell divisions, prophase of meiosis I is extremely prolonged, lasting up to 10 days in male mice (Soh et al., 2017). This process has been histologically described, but a full transcriptional characterization of spermatocytes undergoing meiosis is lacking.

To address this, we ordered spermatocytes along their differentiation trajectory. We first identified a strong increase in the number of genes expressed as spermatocytes progress through prophase, with the highest number being expressed immediately before the cells divide (**Fig. 4A**; **Methods**). Using this as a proxy for active transcription, we identified diplotene spermatocytes, which are the latest cell type in prophase I in which RNA synthesis is occurring (Monesi, 1964). To profile increasing or decreasing transcription throughout meiotic prophase, we correlated each gene’s normalised expression level to the number of genes expressed (**Methods; Table S6**). As expected, previously known marker genes for early meiotic processes such as *Hormad1* and *Sycp3* decreased in expression during Prophase I, whereas *Pou5f2* and *Tcte2*, a male-meiosis specific gene (Braidotti and Barlow, 1997) increased in expression (**Fig. 4B**). Supporting our identification of diplotene spermatocytes, *Pou5f2* has previously been shown to be specifically expressed during a 36-to 48-hour period preceding the meiotic cell division (Andersen et al., 1993).

Despite the overall increase in transcription, we observed distinct temporal expression patterns when visualizing specific marker genes for individual spermatocyte populations. Even within pachytene spermatocytes at different stages in their developmental progression, there exists substantial heterogeneity (**Fig. 4C**). As expected, early spermatocyte markers (SC 1 and SC 2) were enriched for genes with known functions in male or female fertility, reflecting a history of intensive investigation (Deng and Lin, 2002; Spruck et al., 2003; Vries et al., 2005). In contrast, our datasets reveal genes present in previously-unknown transcriptional programs that functionally orchestrate the later stages of spermatocyte development (**Table S2**).

Meiosis culminates in metaphase, where the spindle checkpoint typically eliminates aneuploid cells (Eaker et al., 2001). Whether initiation of the spindle checkpoint results in gene expression perturbation is currently unknown. To explore this, we profiled germ cell development in an aneuploid mouse line (Tc1 mice) that carries one copy of human chromosome 21 (O’Doherty et al., 2005). The aneuploid human chromosome has been associated with frequent congression defects and an arrest at metaphase I; nevertheless, Tc1 mice produce viable sperm, albeit at a reduced level (Ernst et al., 2016).

After profiling single cells throughout spermatogenesis from adult male aneuploid Tc1 mice and matched litter-mate controls (Tc0 mice), we mapped the resulting transcriptomes onto the B6 reference atlas (Methods). As expected, the presence of the human chromosome in cells from the Tc1 mouse resulted in a substantial enrichment in early spermatogenesis immediately prior to meiosis, and post-meiotic cell types were reduced (**Fig. 4D**) (Ernst et al., 2016). Surprisingly, differential expression analysis revealed only a very small number of changes in gene expression on the mouse genome due to the human chromosome (**Fig. S5**). This implies that in the Tc1 mouse model, the sub-fertile phenotype is not driven by transcriptional differences during spermatogenesis. Instead, the cause seems to be the reduced number of cells able to proceed through meiosis in Tc1 mice, with a higher rate of arrest and cell death from activation of the spindle checkpoint, independent of transcription.

In sum, the meiotic gene expression programme is remarkably robust to both aneuploidy as well as variation in the cell types present within tubules.

**Figure 4.**
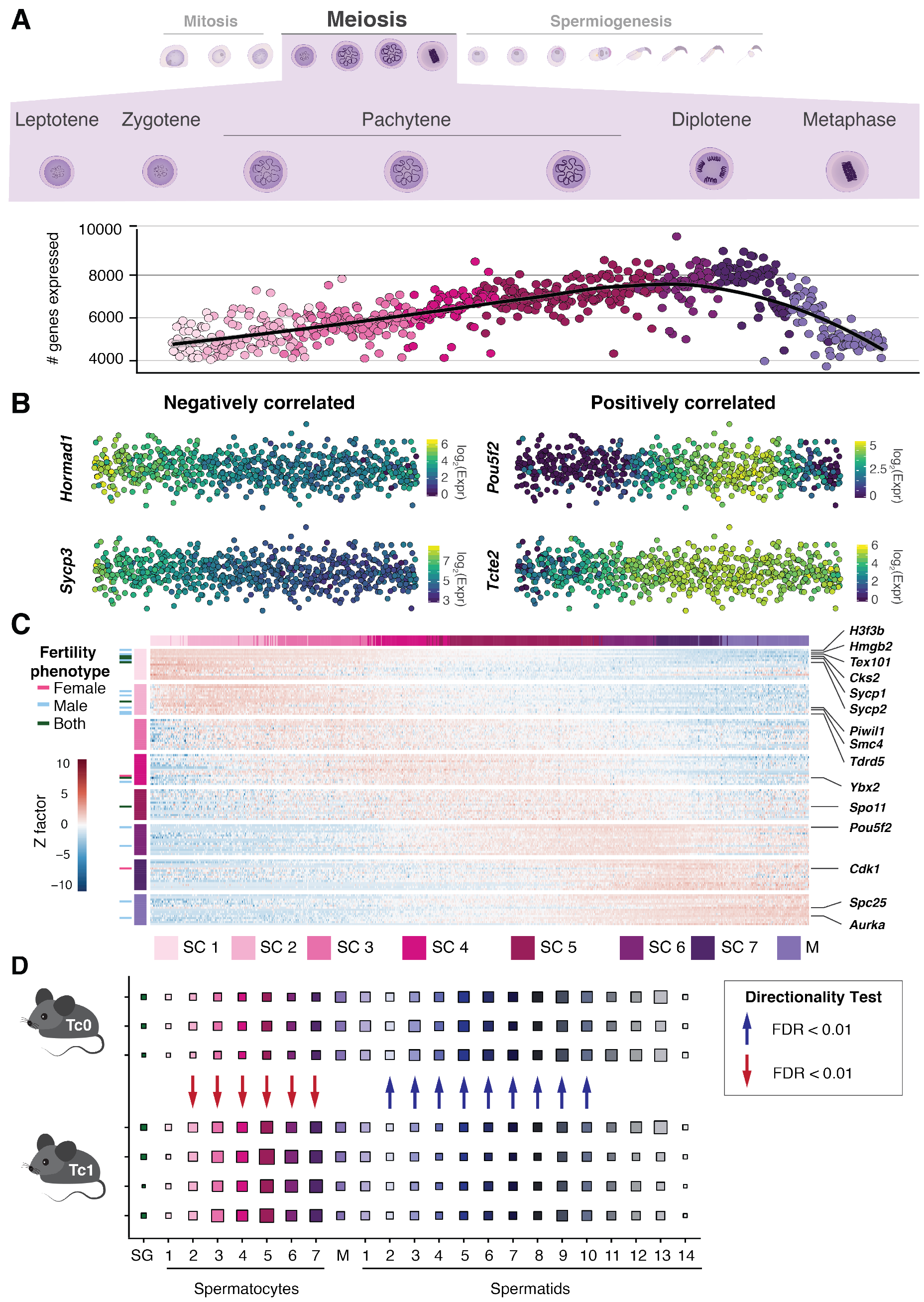
Gene expression dynamics during male meiosis. **(A)** Number of genes expressed per spermatocyte. Cells are ordered by their developmental progression during meiotic prophase until metaphase. **(B)** Expression of genes that are negatively or positively correlated with the number of genes expressed during meiotic prophase (negatively correlated: rho < −0.3, Benjamini-Hochberg corrected empirical p-value < 0.1; positively correlated: rho > 0.3, Benjamini-Hochberg corrected empirical p-value < 0.1, see Methods). Per category, two genes are visualized. The colour gradient represents log_2_-transformed, normalized counts. **(C)** Heatmap visualizing the scaled expression of the top 15 marker genes per cell type. Row and column labels correspond to the different populations of spermatocytes (SC). M: Metaphase. Genes are labelled based on their fertility phenotype: pink - infertile or sub-fertile in females, light blue - infertile or sub-fertile in males, dark green - infertile or sub-fertile in both males and females. **(D)** Cell type proportions for each cluster in Tc0 (n=3) and Tc1 (n=4) animals. Arrows indicate a statistically significant shift in proportions between the genotypes (see Methods). SG - spermatogonia, M - metaphase.

### Transcriptional dynamics of the histone to protamine transition during spermiogenesis

A key event during spermiogenesis is chromatin condensation, which is required to package the haploid genome into the confined space of the sperm nucleus. Our data allowed us to dissect at high-resolution the gradual chromatin remodelling during spermatid differentiation, involving the replacement of canonical histones by histone variants followed by transition proteins and eventually protamines (Balhorn, 2007; El Kennani et al., 2017).

We first explored how expression of histone variants changed throughout early spermatid maturation (**Fig. 5A**). Multiple variants of H3 and H2A are expressed in spermatocytes (Greaves et al., 2006; Mahadevaiah et al., 2001; Tang et al., 2015), and our data showed that many of these histones are highly expressed in early round spermatids. For instance, Histone H3.3 is a histone variant consisting of two genomic copies (*H3f3a* and *H3f3b*). Across spermatogenesis, we observed distinct expression patterns for the two genes, with *H3f3a* being consistently high until the transcriptional shutdown. In contrast, *H3f3b* showed a much more dynamic expression profile, starting high in spermatocytes, dropping throughout meiotic prophase, followed by upregulation in round spermatids (**Fig. S6A**). Although both genes have been implicated in male fertility, the phenotypes associated with perturbations of the more dynamically regulated paralog *H3f3b* are much more severe (Tang et al., 2015; Yuen et al., 2014).

We detected increased expression for particular canonical histones, of which *Hist1h2bp* and *Hist1h4a* showed a distinct up-regulation during early and mid-spermiogenesis (**Fig. S6B**). Canonical histones are typically transcribed in a replication-dependent manner during S phase (Marzluff et al., 2002), thus the atypical expression during spermiogenesis could suggest important roles as replacement histones during chromatin remodelling.

For testis-specific histone variants, we observed highest expression in elongating spermatids, with most variants increasing strongly in expression from S5 onwards. While some variants had a consistently high expression level, *Hils1* and *H1fnt* decreased in expression towards the late stages, similarly to *Tnp1* and *Tnp2* (Zhao et al., 2004). Both histone variants are important for male fertility, and *Hils1* has previously been shown to interact with *Tnp1* (Tanaka et al., 2005; Wu et al., 2009). In contrast, three testis-specific histone variants *Hypm*, *H2afb1* and *H2bl1* (*1700024p04rik*) showed consistently high expression until the end of differentiation similar to protamines, suggesting these variants contribute to the final genome condensation.

As a consequence of chromatin condensation, transcription ceases in spermatids at the round to elongating switch, consistent with the lack of active RNA Pol II at S10 and later stages (Dottermusch-Heidel et al., 2014). Our data clearly reflect this transcriptional shutdown since the number of expressed genes is stable until approximately S9 before gradually declining (**Fig. 5B**).

In the 8 days following transcriptional shutdown, spermatids still need to undergo drastic morphological changes, including the assembly of sperm-specific structures such as the flagellum, before mature testicular sperm can be released into the lumen (O’Donnell, 2015). To achieve this in the absence of active transcription, spermatids store large amounts of mRNAs in a perinuclear RNA granule termed the chromatoid body or *nuage* (Kotaja and Sassone-Corsi, 2007). RNA stored in the chromatoid body is then released for translation, suggesting that these molecules may play vital roles during late stages of spermiogenesis. However, identifying the RNAs that are stored has been hindered by difficulties in purifying late spermatids.

By correlating normalized gene expression against the number of genes expressed, we identified a large number of genes that gradually decrease in relative expression after transcriptional shutdown, the timing of which could be indicative of RNA degradation rates. We reasoned that transcripts where the relative expression after transcriptional shutdown appeared to increase are likely protected from degradation. This included genes with well-known spermiogenesis-specific functions (**Fig 5C**); indeed, transition proteins and protamines are involved in chromatin condensation, as well as genes involved in sperm motility such as *Akap4* and *Cabs1* (Kawashima et al., 2009; Miki et al., 2002). This gene set is therefore a resource for identification of novel spermiogenesis-related proteins with potential roles in fertility (**Table S7**).

**Figure 5:**
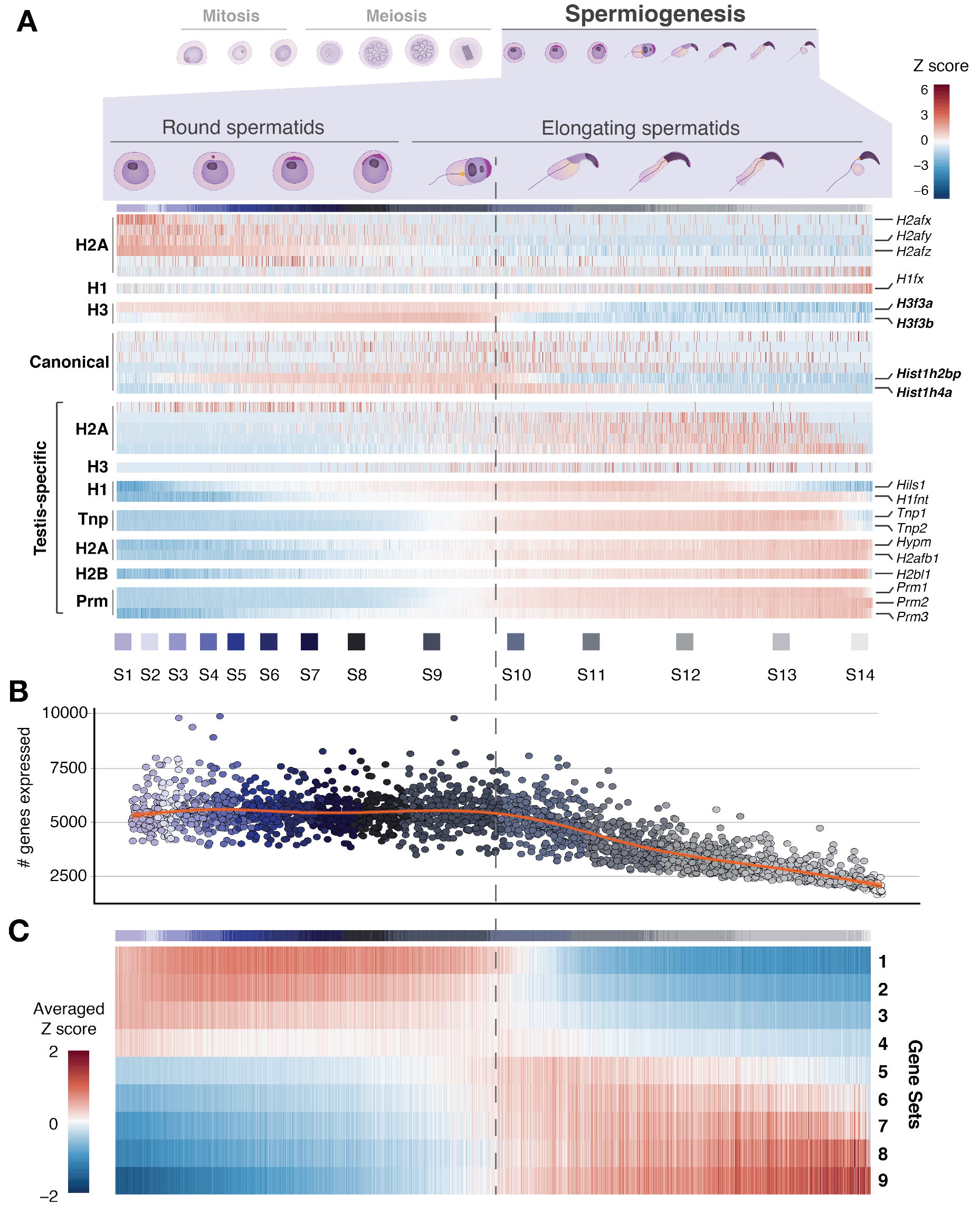
Transcriptional dynamics coupled to chromatin remodelling during spermiogenesis. **(A)** Scaled normalized expression of histone variants (H1, H2A, H2B, H3), canonical histones, transition proteins (Tnp) and protamines (Prm) during spermiogenesis. Cells were ordered based on their developmental trajectory ranging from round spermatids (S1-S8) to elongating spermatids (S9-S14). Vertical dashed line indicates transcriptional shutdown between S9 and S10. **(B)** Number of genes expressed per spermatid. Cells were ordered based on their developmental trajectory. Red line indicates a smooth regression (loess) fit. **(C)** For each gene, its normalised expression per cell was correlated with the number of genes expressed per cell (see Methods). Genes were ordered based on the correlation coefficient and grouped into 9 sets (see Table S7). Scaled expression was averaged across genes within each gene set.

### Meiotic silencing dynamics of sex chromosomes

A male-specific feature of meiosis is the transcriptional silencing of sex chromosomes, followed by partial reactivation in post-meiotic spermatids. This process is termed meiotic sex chromosome inactivation (MSCI), and is caused by asynapsis of the sex chromosomes, leading to accumulation of phosphorylated H2AFX and the formation of the sex body (Hamer et al., 2003) (**Fig. 6A**). By plotting the ratio of gene expression from the X or Y chromosomes compared to all autosomes, the inactivation and re-activation status of the sex chromosomes can be deduced (**Fig. 6B**; **Methods**). The X chromosome is partially upregulated in spermatogonia as described by Sangrithi et al. (2017) (X:A ratio < 1), followed by transcriptional silencing in spermatocytes. Throughout spermiogenesis, expression from the X gradually increases, reaching X:A ratios comparable to spermatogonia, therefore suggesting a substantial reactivation of the X chromosome in post-meiotic spermatids.

Transcriptional silencing was originally thought to persist throughout post-meiotic development (Greaves et al., 2006; Turner et al., 2006); however, several genes have been shown to be re-or *de novo* activated in spermatids, some of which are dependent on Rnf8 (Ring finger protein 8) and/or Scml2 (Sex comb on midleg-like 2) (Hasegawa et al., 2015; Sin et al., 2012, 2015). However, the precise timing and order of the transcriptional reactivation of *de novo* escape genes during spermiogenesis has not been explored.

Profiling whole-testis transcriptomes of juvenile mice sampled every two days during the first wave of spermatogenesis allowed the sensitive detection of spermatid-specific escape genes (**Fig. 2C**). Due to the gradual emergence of germ cell types during the first spermatogenic wave, differential expression analysis between early (< P20) and late (> P20) time points revealed genes exclusively expressed in spermatids and which are thus *de novo* activated escape genes (n = 128) (**Fig. 6C**; **Table S8**). These include many of the previously annotated escape genes such as *Cypt1*, *Cycl1*, and *Akap4*, and show an enrichment for genes dependent on Rnf8 or Scml2 for reactivation in spermatids (Fisher’s Exact Test: RNF8-targets, p-value < 5×10^−12^; Scml2-targets, p-value < 2×10^−9^) (**Fig. 6C**) (Adams et al., 2018).

The *de novo* activated genes across our single cell RNA-Seq dataset showed a broad range of temporal expression patterns (**Fig. 6D**). The earliest expression, directly following meiosis and lasting until stages S4-S5 was observed for three members of the *Ssxb* multicopy gene family (*Ssxb1*, *Ssxb2*, *Ssxb3*). Multi-copy genes have previously been described to have spermatid-specific expression (Mueller et al., 2008), and their ampliconic structure has been speculated to play a role in escaping meiotic silencing *via* self-pairing (Disteche, 2008). Other multi-copy genes showing a spermatid-specific expression pattern included *Rhox11*, *Mageb5* and *Slxl1*. However, no other multi-copy gene families showed early reactivation similar to *Ssxb*, which suggests that the *Ssxb* gene family may have distinct function in X reactivation post meiosis (**Fig. S7**).

### Epigenetic mechanisms underlying *de novo* escape gene activation

To gain insight into the mechanisms underlying *de novo* activation of spermatid-specific escape genes on the X chromosome, we profiled the chromatin landscape in spermatocytes and spermatids using the newly developed CUT&RUN protocol for low cell numbers (**Fig. 7A**; **Fig. S8A; Methods**, (Skene et al., 2018)). We assayed trimethylation of histone H3 on lysine 4 (H3K4me3) as a proxy for promoter activity, as well as repressive trimethylation of lysine 9 (H3K9me3), which is associated with the sex body in early and late spermatocytes and enriched in post-meiotic sex chromatin (PMSC) (Greaves et al., 2006; Tachibana et al., 2007). By profiling the enrichment of H3K9me3 across all chromosomes, we confirmed that the X chromosome has high levels of H3K9me3 in spermatids (Moretti 2016 Epigenetics Chromatin). In addition, we now show that H3K9me3 accumulation begins earlier in meiosis, and indeed spermatocytes show enrichment of this repressive mark (**Fig. 8B**).

On autosomes, H3K9me3 is enriched in pericentromeric regions of constitutive heterochromatin (Peters et al., 2001); in sharp contrast, this repressive histone mark is more evenly distributed across the X chromosome in spermatocytes (**Fig. S8C**). Nevertheless, we detected broad regions showing particularly high levels of H3K9me3 scattered across the X chromosome (**Fig. S8B**), including the promoter of *Akap4*, a well-known escape gene. This discovery prompted us to profile the chromatin dynamics of active and repressive marks at promoters of *de novo* escape genes (*spermatid-specific genes*) versus the promoters of all other expressed X-chromosome genes (*non-spermatid specific genes*) (**Fig. 6C**; **Table S8**).

In spermatocytes, spermatid-specific genes showed lower enrichment in H3K4me3 than non-spermatid specific genes (Wilcoxon-Mann-Whitney: p-value < 2.2×10^−16^) (**Fig. 7C**, left panel). In contrast, spermatid-specific genes have on average elevated H3K4me3 in spermatids, as expected based on their increased expression level compared to spermatocytes (**Fig. 7C**, right panel). When examining the deposition of H3K9me3 on the promoters of X-linked genes, we detected a strong enrichment in spermatid-specific escape genes in spermatocytes (Wilcoxon-Mann-Whitney: p-value < 3.7×10^−11^) (**Fig. 7D**). This pattern indicates that spermatid-specific genes are more strongly repressed in spermatocytes.

Our results describe for the first time, the precise epigenetic changes associated with escape gene activation in post-meiotic cells. These dynamics are exemplified by the chromatin remodelling that occurs around *Akap4* and *Cypt1*, both of which are well-studied spermatid-specific genes (**Fig. 7E**). The promoters of these genes have high levels of H3K9me3 in spermatocytes, which decreases in spermatids, while H3K4me3 levels are strongly increased.

**Figure 6:**
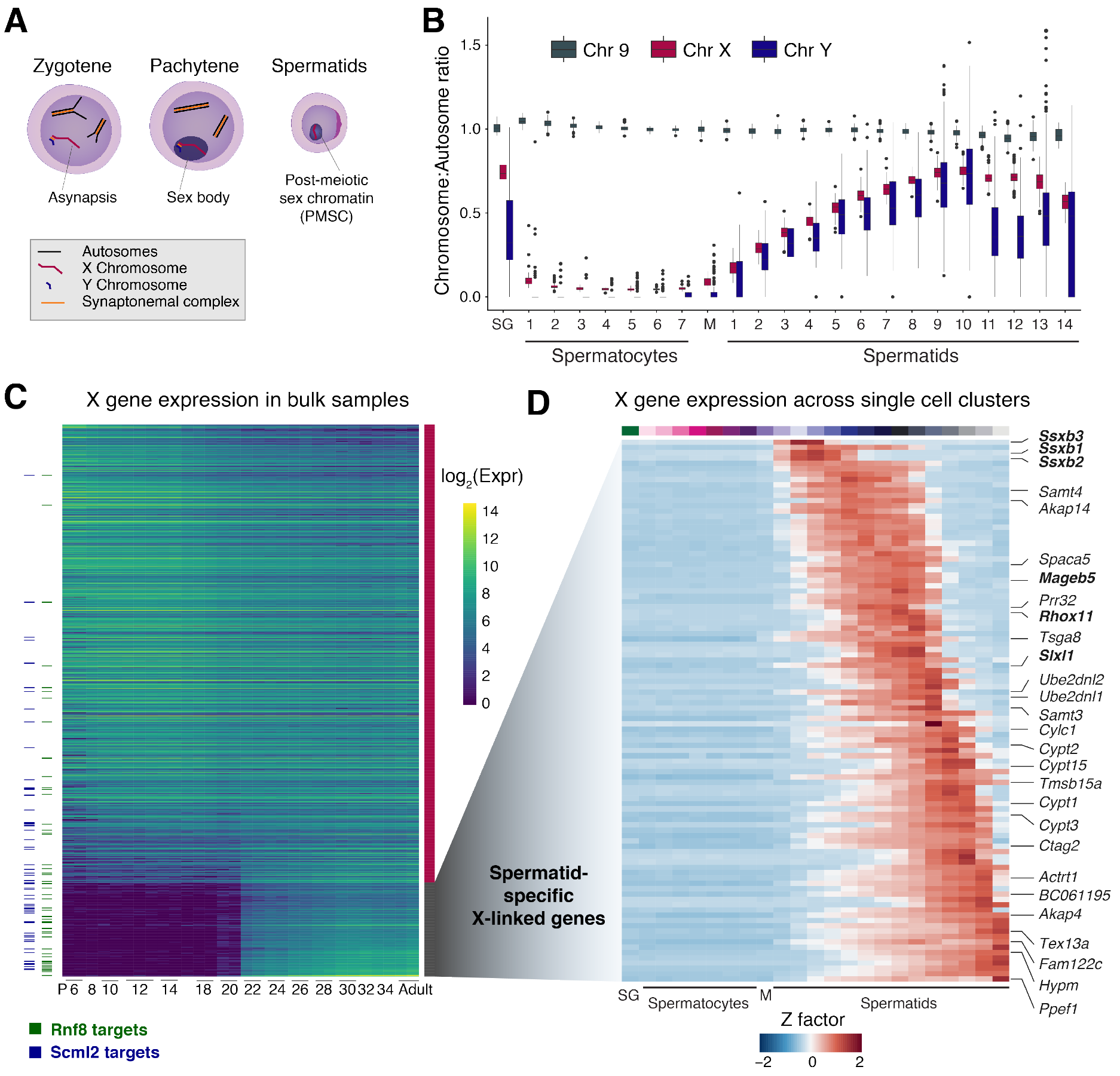
X chromosome dynamics during spermatogenesis. **(A)** Schematic of sex chromosome sub-nuclear localisation through spermatogenesis. **(B)** For each cell, the ratio of mean expression of genes on Chr 9, Chr X and Chr Y to the mean expression of genes across all autosomes is represented as a boxplot for cells allocated to each developmental stage (see Methods). SG - spermatogonia, M - metaphase. **(C)** Expression of all X chromosome genes (> 10 average counts) in bulk RNA-seq data across the juvenile time course. Columns correspond to developmental stage and rows are ordered by the log_2_ fold change between spermatocytes (stages before postnatal day (P) 20) and spermatids (stages after and including P20). Horizontal dashes indicate genes that are targets of Rnf8 (green) and Scml2 (blue) (Adams et al., 2018). **(D)** Average expression of spermatid specific genes (panel (C)) per germ cell type. Columns are ordered by developmental stage and rows are ordered by peak gene expression through development. Multi-copy genes are highlighted in bold.

**Figure 7:**
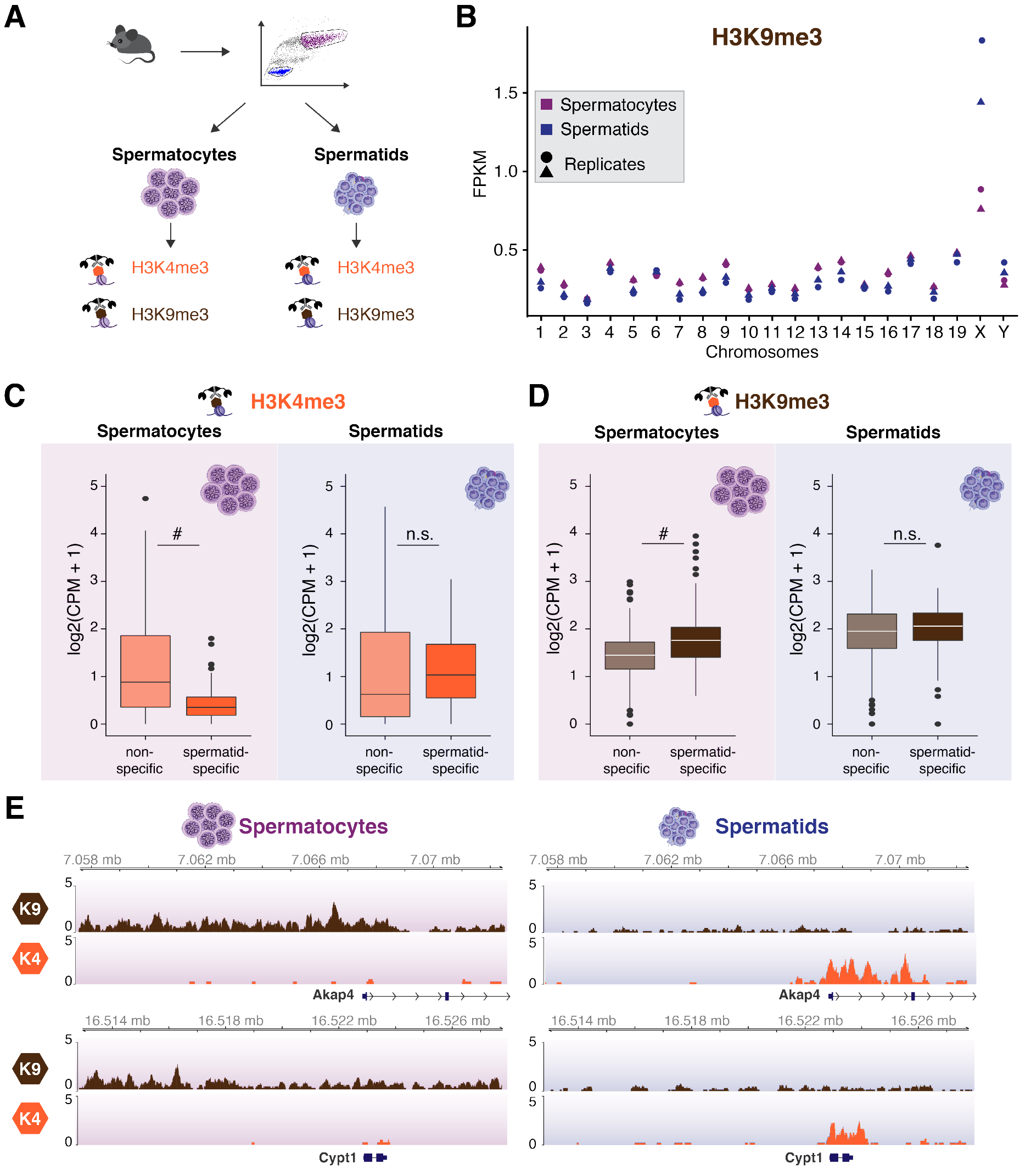
Profiling of repressive and active chromatin marks in spermatocytes and spermatids. **(A)** Spermatocytes and spermatids were isolated from the same individual using FACS and profiled using H3K4me3 (active mark) and H3K9me3 (repressive mark) using CUT&RUN. **(B)** Number of H3K9me3 Fragments Per Kilobase per Million (FPKM) for each chromosome. Pink (spermatocytes); Blue (spermatids). Shape corresponds to biological replicate. **(C)** and **(D)** Boxplot of H3K4me3 **(C)** and H3K9me3 **(D)** Counts Per Million (CPM) in promoter regions of spermatid specific (n=127) and non-spermatid specific (n=617) genes for spermatocytes (left) and spermatids (right). # indicates statistical significance (Wilcoxon-Mann-Whitney: p-value < 1×10^−10^), n.s. - not significant. **(E)** Genome tracks of H3K4me3 and H3K9me3 for two representative spermatid-specific genes (*Akap4* and *Cypt1*) for Spermatocytes (left) and Spermatids (right). Reads were scaled by library size.

## DISCUSSION

The testes are among the most proliferative tissues in the adult body and ensure fertility via the continuous production of millions of sperm per day. In contrast to most developmental differentiation processes which require the profiling of cellular populations at several time points (Kernfeld et al., 2018; Scialdone et al., 2016; Wagner et al., 2018), spermatogenesis occurs in continuous waves throughout the reproductive life span, with all intermediate cell types that arise across the ~35 day differentiation program present in adult testes. This provided a powerful opportunity to capture and profile an entire differentiation process by profiling the transcriptomes of thousands of single-cells at a single time point.

We identified key developmental transitions within the differentiation trajectory by profiling the first wave of spermatogenesis. Because germ cells have only progressed to a defined developmental point, the differentiation trajectory was truncated, facilitating identification of the most mature cell type. Profiling spermatogenesis in juvenile animals also naturally enriched for rare cell-types that are under-represented in adults. Among these, spermatogonia are of particular interest as these cells not only sustain male fertility, but are also the origin of the vast majority of testicular neoplasms (Bosl and Motzer, 1997). We obtained more than 1100 transcriptional profiles for spermatogonia, allowing the identification of specific cell clusters within this heterogeneous cell population thus greatly improving the resolution over previous studies that only studied adult testes (Lukassen et al., 2018). Furthermore, our approach also enriched for and facilitated characterisation of the complexity within testicular somatic cell types, thus providing a valuable resource for understanding tissue homeostasis.

Droplet-based scRNA-Seq can profile large number of cells simultaneously (Klein et al., 2015; Macosko et al., 2015; Zheng et al., 2017), but often captures cells with a wide range of transcriptional complexity. Consequently, droplet-based assays present a major computational challenge in distinguishing between (i) droplets contain transcriptionally inactive cells versus (ii) empty droplets that contain (background) ambient RNA. By using a stringent default threshold, we identified the majority of somatic and germ cell types in testes, similar to recent single-cell expression studies in mouse and human (Lukassen et al., 2018; Xia et al., 2018). In addition, we applied a new statistical method to identify cells from droplet-based data by comparing the ambient RNA profiles (Lun et al., 2018), and were able to identify transcriptionally inactive leptotene/zygotene spermatocytes. This allowed us to bridge the developmental transition between spermatogonia and spermatocytes, thus providing a more complete view of the continuum of germ cell differentiation.

Perturbations of the gene expression programme during spermatogenesis frequently result in male sterility by causing a maturation arrest (Cooke and Saunders, 2002). It is therefore of great interest to identify genes with novel spermatogenesis-related functions. To this end, we further characterized the dynamic gene expression patterns underlying meiosis and spermiogenesis. Although transcription broadly increases during meiotic prophase, we detected a diverse set of regulatory behaviours, particularly among spermatocyte-specific genes.

In the late stages of spermiogenesis, transcription ceases due to the histone-to-protamine transition. Remarkably, the transcripts from a large set of genes remain highly abundant even after transcriptional shutdown in elongating spermatids. Many of these genes are known to be essential for reproductive success and therefore our analysis likely reveals novel genes with roles in sperm maturation.

The transcriptional silencing of the sex chromosomes during meiosis and their subsequent partial re-activation post-meiosis is essential for male fertility (Mahadevaiah et al., 2008). Failure of meiotic sex chromosome inactivation (MSCI) results in the expression of spermatocyte-lethal genes, as demonstrated for two Y chromosome encoded genes zinc finger protein Y-linked (*Zfy*) 1 and 2 (Royo et al., 2010). Our discovery that H3K9me3 is enriched during meiosis at spermatid-specific genes suggests a stronger, targeted repression in spermatocytes for a key subset of X-linked genes. The deposition of H3K9me3 is specific to MSCI in males, and is not observed during general meiotic silencing of unpaired chromosomes (MSUC) (Cloutier et al., 2016; Taketo and Naumova, 2013; Turner et al., 2004b). Interestingly, when comparing the levels of meiotic silencing for the X chromosome in spermatocytes with the unpaired X chromosome in *XO* oocytes, the transcriptional silencing was stronger in males compared to females (Cloutier et al., 2016). The deposition of H3K9me3 is linked to more robust silencing of X-chromosomal genes; our finding that spermatid-specific genes are particularly enriched for H3K9me3 in spermatocytes suggests that their repression may be necessary for male fertility.

Such a requirement could arise from the opposing evolutionary forces acting on the X chromosome (Rice, 1984, 1992). Due to its hemizygosity in males, the X chromosome has been predicted to be enriched for male-specific genes. In contrast, meiotic silencing allows pachytene-lethal genes to survive on the X chromosome, since their deleterious effect will be masked by MSCI, similarly to *Zfy1/2* on the Y chromosome (Royo et al., 2010). Our study thus raises interesting questions about how H3K9me3 is targeted to specific genes on the X chromosome in spermatocytes, and how transcription is reactivated in post-meiotic spermatids.

## MATERIALS AND METHODS

### Mouse material

All animals were housed in the Biological Resources Unit (BRU) in the Cancer Research UK - Cambridge Institute under Home Office Licences PPL 70/7535 until February 2018 and PPL P9855D13B from March 2018. C57BL/6J animals were purchased from Charles River UK Ltd (Margate, United Kingdom) and the Tc1 mouse line was obtained from Dr. E. Fisher and Dr. V. Tybulewizc (O’Doherty et al., 2005) and maintained by breeding female Tc1 mice to male (129S8 x C57BL/6J) F1 mice. Littermates that did not inherit human chromosome 21 in these crosses (Tc0) were used as control animals.

### Fluorescence-activated cell sorting of spermatogenic cell populations

Spermatogenic cell populations were isolated from adult mouse testes as described in Ernst et al. (2016). In brief, the albuginea was removed and tissue was incubated in dissociation buffer containing 25 mg/ml Collagenase A, 25 mg/ml Dispase II and 2.5 mg/ml DNase I for 30 minutes at 37°C. Enzymatic digestion was quenched with Dulbecco’s Modified Eagle Medium (DMEM, Gibco) supplemented with 10% Fetal calf serum (FCS, 10270106, Gibco). Cells were resuspended at a concentration of 1 million cells per ml and stained with Hoechst 33342 (H3570, ThermoFisher Scientific) at a final concentration of 5 *μ*g/ml for 45 minutes at 37°C. Cells were resuspended in PBS containing 1% FCS and 2 mM EDTA and propidium iodide was added to a final concentration of 1 *μ*g/ml prior to sorting.

Cells were sorted on an Aria IIu cell sorter (Becton Dickinson) using a 100 *μ*m nozzle. Hoechst was excited with a UV laser at 355nm and fluorescence was recorded with a 450/50 filter (Hoechst blue) and 635LP filter (Hoechst red). Primary spermatocytes (4N) and round spermatids (1N) were sorted and collected in PBS containing 1% FCS and 2 mM EDTA.

### Total RNA-Seq from bulk samples

Testes from prepubertal mice ranging between postnatal day 6 and 35 were flash frozen or directly used for RNA extraction using Trizol (Thermo Fisher, 15596026) following manufacturer’s instructions. Purified RNA was DNase-treated using the TURBO DNA-free Kit according to manufacturer’s instructions (Thermo Fisher, AM1907) and RNA quality was assessed using the Agilent Tapestation RNA Screentape. 800 ng of DNA-depleted RNA were used for RNA-Seq library preparation using the TruSeq Stranded Total RNA Library Kit with Ribo-Zero Gold for cytoplasmic and mitochondrial ribosomal RNA removal according to manufacturer’s instructions (Illumina, RS-122-2303). Libraries were then sequenced on Illumina HiSeq2500 using a paired-end 125bp run.

### 10X Genomics Single-cell RNA-Seq

Mouse testes were enzymatically dissociated as described above and 34 *μ*l of singlecell suspension at a concentration of ~297,000 cells/ml was loaded into one channel of the Chromium™ Single Cell A Chip (10X Genomics^®^), aiming for a recovery of 4000-5000 cells. The Chromium Single Cell 3’ Library & Gel Bead Kit v2 (10X Genomics^®^, 120237) was used for single-cell barcoding, cDNA synthesis and library preparation, following manufacturer’s instructions according to the Single Cell 3’ Reagent Kits User Guide Version 2, Revision D. Libraries were sequenced on Illumina HiSeq2500 using a paired-end run sequencing 26bp on read 1 and 98bp on read 2. Information about libraries in which individual samples were sequenced is available in **Table S1**.

### Histology

Testes were fixed in neutral buffered formalin (NBF) for 24 hours, transferred to 70% ethanol, machine processed and paraffin embedded. Formalin-fixed paraffin-embedded (FFPE) sections of 3um thickness were used for all histological stains and immunohistochemistry (IHC).

For Periodic Acid Schiff (PAS) stainings slides were dewaxed, washed in water and placed in 0.5% Periodic Acid (Sigma P0430) for 5 minutes. After three washes in ultra-pure water, slides were placed in Schiff reagent (Thermo Fisher Scientific, J/7300/PB08) for 15-30 minutes in a closed container and washed again three times in ultra-pure water. Counterstain was performed using Mayers Haematoxylin (Thermo Fisher Scientific, LAMB/170-D) for 40 seconds followed by rinsing in tap water, dehydration and mounting.

IHC was performed on FFPE sections using the Bond™ Polymer Refine Kit (DS9800, Leica Microsystems) on the automated Bond Platform. Anti-phospho-Histone H3 (Ser10) (pH3) antibody (Upstate, 06-570, 1:200 dilution) was used with DAB Enhancer (Leica Microsystems, AR9432) and heat-induced epitope retrieval was performed for 10 minutes at 100°C on the Bond platform with sodium citrate. All slides were scanned using Aperio XT (Leica Biosystems) and PH3 intensities were quantified using the Aperio eSlide Manager (Leica Biosystems).

### Low cell number chromatin profiling using CUT&RUN (Cleavage under targets and release using nuclease)

*In situ* chromatin profiling of FACS-purified spermatogenic cell populations was performed according to Skene et al. (2018) In brief, spermatocytes and spermatids were sorted as described above and collected in PBS. Cells were spun down at 600 g for 3 minutes in swinging-bucket rotor and washed twice with 1.5 ml Wash buffer (20 mM HEPES-KOH (pH 7.5), 150 mM NaCl, 0.5 mM Spermidine and 1X cOmplete™ EDTA-free protease inhibitor cocktail (04693159001, Roche)). During the cell washes, concanavalin A-coated magnetic beads (Bangs Laboratories, cat. No BP531) (10 *μ*l per condition) were washed twice in 1.5 mL Binding Buffer (20 mM HEPES-KOH (pH 7.5), 10 mM KCl, 1mM CaCl, 1mM MnCl_2_) and resuspended in 10 ul Binding Buffer per condition. Cells were then mixed with beads and rotated for 10 minutes at room temperature (RT) and samples were split into aliquots according to number of antibodies profiled per cell type. We used 20,000-30,000 spermatocytes and 40,000-60,000 spermatids per chromatin mark.

Cells were then collected on magnetic beads and resuspended in 50 *μ*l Antibody Buffer (Wash buffer with 0.05% Digitonin and 2 mM EDTA) containing one of the following antibodies in 1:100 dilution: H3K4me3 (Millipore 05-1339 CMA304, Lot2780484) and H3K9me3 (Abcam, ab8898, Lot GR306402-1). Cells were incubated with antibodies for 10 minutes at RT and then washed once with 1 ml Digitonin buffer (Wash buffer with 0.05% Digitonin). For the mouse anti-H3K4me3 antibody, samples were incubated with a 1:100 dilution in Digitonin buffer of secondary rabbit anti-mouse antibody (Invitrogen, A27033, Lot RG240909) for 10 minutes at RT and then washed once with 1 mL Digitonin buffer. Samples were then incubated with 700 ng/ml ProteinA-MNase fusion protein (kindly provided by Steven Henikoff) for 10 minutes at room temperature followed by two washes with 1 ml Digitonin buffer. Cells were then resuspended in 100 *μ*l Digitonin buffer and cooled down to 4°C before addition of CaCl_2_ to a final concentration of 2 mM. Targeted digestion was performed for 30 minutes on ice until 100 *μ*l of 2X STOP buffer (340 mM NaCl, 20 mM EDTA, 4 mM EGTA, 0.02% Digitonin, 250 mg RNase A, 250 *μ*g Glycogen, 15 pg/ml yeast spike-in DNA (kindly provided by Steven Henikoff)) were added. Cells were then incubated at 37°C for 10 minutes to release cleaved chromatin fragments, spun down for 5 minutes at 16,000 g at 4°C and collected on magnet. Supernatant containing the cleaved chromatin fragments was then transferred and cleaned up using the Zymo Clean & Concentrator Kit.

Library preparation was performed using the ThruPLEX^®^ DNA-Seq Library Preparation Kit (R400407, Rubicon Genomics) with a modified Library Amplification programme: Extension and cleavage for 3 minutes at 72°C followed by 2 minutes at 85°C, denaturation for 2 minutes at 98°C followed by four cycles of 20 seconds at 98°C, 20 seconds at 67°C and 40 seconds at 72°C for the addition of indexes. Amplification was then performed for 12-14 cycles of 20 seconds at 98C and 15 seconds at 72C. Average library size was tested on Agilent 4200 Tapestation using a DNA1000 High Sensitivity Screentape and quantification was performed using the KAPA Library Quantification Kit (Kapa Biosystems). CUT&RUN libraries were sequenced on a HiSeq2500 using a paired-end 125bp run.

### Processing of 10X Genomics scRNA-Seq data

#### Read alignment and counting of 10X Genomics scRNA-Seq data

To generate a genomic reference for sequence alignment, the full *Mus musculus* genome (GRCm38) was concatenated with the sequence of the human chromosome 21 (taken from GRCh38). Similarly, the genomic annotation for *Mus musculus* (GRCm38.88) was merged with the annotation for human chromosome 21 (taken from GRCh38.88). The Cell Ranger v1.3.1 *mkref* function with default settings was used to process the genomic sequence and the annotation file for read alignment. To obtain gene-specific transcript counts, the Cell Ranger v1.3.1 *count* function with default settings was used to align and count unique molecular identifiers (UMIs) per sample.

#### Quality control of Cell Ranger filtered cells

The Cell Ranger v1.3.1 software retains cells with similar UMI distributions (Zheng et al., 2017). We use this default threshold to obtain high-quality cells with large numbers of UMIs. After merging all samples, we filtered out cells that express less than 1000 genes. Furthermore, we exclude cells with more than 10% of reads mapping to the mitochondrial genome. These filtered data were used for all analyses except that presented in Figure S4 and Table S5 where the *emptyDrops* filtered cells (below) were utilised.

#### Quality control of emptyDrops filtered cells

Using the Cell Ranger default threshold leads to the exclusion of cells with lower transcriptional complexity. We therefore used the *emptyDrops* function provided in the *DropletUtils* Bioconductor package (Lun et al., 2018) to statistically distinguish empty droplets from genuine cells (controlling the FDR to 1%). After merging true cells across all samples, we filtered out cells with less than 500 genes expressed. Furthermore, we excluded cells with more than 10% or mitochondrial genes expressed.

#### Normalization

The transcriptomes of quality filtered cells were normalized using the *scran* package (Lun et al., 2016). Cells with similar transcriptomic complexity were pre-clustered using a graph-based approach (as implemented in the *quickCluster* function). Size factors were calculated within each cluster before being scaled between clusters using the *computeSumFactors* function. Throughout this paper, the log_2_-transformed, normalized counts (after adding one pseudocount) are displayed. For down-stream analysis, lowly expressed genes (averaged log_2_-transformed, normalized expression < 0.1) were excluded.

#### Detection of highly variable genes

To detect the top 1000 most variable genes across all tested cells, we first fitted a regression trend between the variance of the log_2_-transformed normalized counts and the abundance of each endogenous gene using the *trendVar* function in *scran*. Next, we used the *decomposeVar* function in *scran* to compute the biological variation for each gene (Lun et al., 2016b). Genes are ordered based on their biological variation and the top 1000 most variable genes are selected.

#### Computational mapping of single cells across samples

To remove batch-specific effects that arise when samples are prepared and sequenced in different experiments (Table S1), we used the *mnnCorrect* function implemented in the *scran* package (Haghverdi et al., 2018). We used the top 1000 genes with highest biological variation across all samples as informative genes for batch correction. Batch correction was performed across (i) all samples, (ii) P10 and P15 spermatogonia or (iii) P10 and P15 spermatogonia and early spermatocytes using *mnnCorrect* with the following parameters: cos.norm.in=TRUE, cos.norm.out=TRUE, sigma=0.1. The *mnnCorrect* function takes transcriptional profiles of cells isolated from adult B6 mice as first input and uses this dataset as reference for cell mapping.

#### Clustering of batch-corrected single-cell transcriptomes

The batch corrected transcriptomes (using the shared top 1000 highly variable genes as explained above) were clustered using a graph-based approach. A shared nearest-neighbour (SNN) graph (Xu and Su, 2015) was constructed considering 3 shared nearest neighbours using the *buildSNNGraph* function in *scran*. In the next step, a multi-level modularity optimization algorithm was used to find community structure in the graph (Blondel et al., 2008) implemented in the *cluster_louvain* function of the *igraph* R package. Clusters were annotated based on the expression of known markers genes. Cells in small clusters that show unclear identities were excluded from down-stream analysis.

Clustering of the P15 sample after emptyDrops filtering was performed on the log_2_-transformed, normalized counts using 10 shared nearest neighbours and the same strategy as explained above. Clustering of the batch-corrected counts of P10 and P15 spermatogonia was performed using 15 shared nearest neighbours.

#### Dimensionality reduction and hierarchical clustering

For visualization, t-distributed stochastic neighbour embedding (tSNE) was computed on the batch-corrected counts of all samples using the R package *Rtsne*. Throughout this study, we visualize subsets of this tSNE. Principal component analysis (PCA) was computed either on the log_2_-transformed, normalized counts of the top 1000 most highly variable genes or batch-corrected counts of the scRNA-Seq data using the base R *prcomp* function. To visualise the cell-to-cell distances of their transcriptomes, we performed hierarchical clustering on log_2_-transformed, normalized counts of the top 1000 most highly variable genes using the base R *hclust* function. We used the Spearman correlation (van Dongen and Enright, 2012) as a distance metric and ward.D2 as clustering strategy.

#### Differential expression testing and marker gene extraction

Differential expression testing across multiple pairwise comparisons was used to identify cluster-specific marker genes in the adult B6 samples, juvenile P10 samples, spermatogonia of P10 and P15 animals and cells detected in P15 sample after emptyDrops filtering. To detect cluster-specific marker genes, the *findMarkers* function implemented in *scran* was used on the log_2_-transformed normalized counts while providing the cluster labels.

To detect differentially expressed genes between Tc1 and Tc0 animals, we summed counts within each cell cluster and each batch to form *pseudo-bulk* samples. We used the Bioconductor package *edgeR* (*Robinson et al., 2010)* to perform differential expression analysis between the pseudo-bulk samples of Tc0 and Tc1 (testing an absolute log-fold change in mean expression > 0.5). This approach avoids confounding batch effects between the two genotypes (Lun and Marioni, 2017).

#### Differential cell-proportion testing between samples

To statistically test for differences in cell type proportions between Tc0 and Tc1 animals, we counted the number of cells allocated to each cell type within each batch. *EdgeR* was used to perform differential proportion testing by scaling the number of cells per cell type by the total number of cells per batch.

#### Ordering cells along their developmental trajectory

To order cells along their developmental trajectory, we fitted a principal curve (Hastie and Stuetzle, 1989) to the first 3 principal components (computed on the top 1000 highly variable genes) using the *principal.curve* function implemented in the *princurve* R package. This approach allows us to order cells along the principal curve. The directionality of the curve was inferred using prior information based on the cluster annotation.

#### Correlation analysis

To correlate log_2_-transformed normalized gene expression to the number of genes expressed, we used the *correlatedPairs* function implemented in *scran (Lun et al., 2016b)*. We first constructed an empirical null distribution using the *correlateNull* function implemented in *scran*. Next, we tested whether the observed Spearman’s rho for each gene against this null distribution. We consider genes with rho < −0.3 and a Benjamini-Hochberg corrected empirical p-value < 0.1 as negatively correlated and genes with rho > 0.3 and a Benjamini-Hochberg corrected empirical p-value < 0.1 as positively correlated.

#### Computing the sex chromosome to autosome ratio

To compute the ratio in expression between chromosome 9, chromosome X or chromosome Y and all autosomes, we selected genes that were expressed in more than 30% of spermatogonia or 30% of spermatids, the cell types with detectable sex chromosome expression. For each cell, the mean expression across these genes per chromosome was calculated. Mean expression of the chromosomes of interest (9, X and Y) was divided by mean expression of the autosomes.

### Processing of bulk RNA-Seq data

#### Read alignment and counting of bulk RNA-Seq data

Sequencing reads were aligned against the *Mus musculus* genome (GRCm38) using the STAR aligner v2.5.3 (Dobin et al., 2013) with default settings. Gene-level transcript counts were obtained using HTSeq version 0.9.1 (Anders et al., 2015) with the −s option set to “reverse” and using the GRCm38.88 genomic annotation file.

#### Quality control and normalization

We visualized several features of the aligned and counted data (number of intronic/exonic reads, number of multi-mapping reads, low-quality reads and total library size) and did not detect any low-quality RNA-Seq libraries. Next, we used the size factor normalization approach implemented in *DESeq2* (Love et al., 2014) for data normalization. For down-stream analysis and visualization, lowly expressed genes (averaged counts < 10) were excluded.

#### Probabilistic classification of bulk samples

We used a regression approach to link the bulk samples to the transcriptomic profiles of single cells. Using the top 50 cluster-specific marker genes for spermatogonia, all spermatocyte groups, all spermatid groups, sertoli and leydig cells, we trained a random forest classifier (implemented in the *randomForest* R package (Liaw and Wiener, 2002)) on 2000 cells isolated from adult B6 testes. Model testing was performed on the remaining 1215 cells isolated from adult B6 testes. Prior to training and testing, log_2_-transformed, normalized counts were scaled by computing the Z score for each gene. Probabilistic prediction was performed using the Z score of log_2_-transformed, normalized bulk RNA-Seq reads of the input genes.

#### Differential expression analysis between spermatocytes and spermatids

Differential expression analysis between cells present before post-natal day 20 and after day 20 was performed using *edgeR*. The *glmTreat* function was used for testing with a minimum absolute log_2_-fold change threshold > 2. Spermatid-specific genes are identified with a log_2_-fold change > 5 in samples after day 20 compared to samples before day 20 (controlling the FDR to 10%).

### Processing of CUT&RUN data

#### Read alignment of CUT&RUN data

Reads were aligned to the *Mus musculus* genome (GRCm38) using Bowtie2 with the following settings: --local --very-sensitive-local --no-unal-q --phred33.

#### Read counting in specified regions

Paired end reads were counted in specified regions using the *regionCounts* function implemented in the *csaw* Bioconductor package (Lun and Smyth, 2016). For this, duplicated reads, reads mapped more than 1000 bp apart and reads mapping to blacklisted regions (available at: http://mitra.stanford.edu/kundaje/akundaje/release/blacklists/mm10-mouse/mm10.blacklist.bed.gz) were removed. Regions of interests were: promoters (obtained using the *promoters* function of the *GenomicFeatures* package), 1000 bp windows across the chromosome (using the *windowCounts* function of *csaw*) and whole chromosomes.

#### Scale normalization of counted reads

Counts per region were normalized based on library size (counts per million, CPM) for promoter regions and 1000 bp windows; additionally, when considering entire chromosomes, the length of the chromosome was accounted for by computing the Fragments per Kilobase per Million mapped reads (FPKMs).

#### Regions with high H3K9me3 counts

To visualize regions with the highest H3K9me3 signal, we merged the top 1000 windows (1000 bp width) using the *mergeWindows* of the *csaw* package with a tolerance of 1500 bp.

### Gene annotation

We performed literature search and used the database www.mousephenotype.org to annotated genes regarding their sterility phenotype. To visualize histone variants and canonical histones, we used the annotation found in El Kennani et al. (2017). Targets of Rnf8 and Scml2 were taken from Adams et al. (2018)

### Multi-copy gene analysis

To analyse multi-copy gene families, we used the annotation from Mueller et al. (2008), Table S1. The cDNA sequence of these genes was obtained from Ensembl (www.ensembl.org) and the BLAST tool was used to identify sequences of genes with high similarity (> 90%). For each multi-copy gene family, normalized counts for all X-chromosomal genes with high sequence similarity were summed.

### Visualization of expression counts

To visualize gene-level transcript counts we either plot the log_2_-transformed normalized counts or the Z score of the log_2_-transformed, normalized transcript counts. The Z score is computed as: 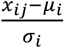 where x is the log_2_-transformed, normalized count for gene i in cell j, *μ*_*i*_ is the mean of gene i across all cells and *σ*_*i*_ is the standard deviation for gene i across all cells.

### Ethics statement

This investigation was approved by the Animal Welfare and Ethics Review Board and followed the Cambridge Institute guidelines for the use of animals in experimental studies under Home Office licences PPL 70/7535 until February 2018 and PPL P9855D13B from March 2018. All animal experimentation was carried out in accordance with the Animals (Scientific Procedures) Act 1986 (United Kingdom) and conformed to the Animal Research: Reporting of *In Vivo* Experiments (ARRIVE) guidelines developed by the National Centre for the Replacement, Refinement and Reduction of Animals in research (NC3Rs).

## Data availability

Data have been deposited under ArrayExpress accession number E-MTAB-6946 for scRNA-Seq data, E-MTAB-6934 for bulk RNA-Seq data and E-MTAB-6932 for CUT&RUN data and are currently under curation.

## Code availability

The R code to reproduce the full analysis and all figures can be obtained from: https://github.com/MarioniLab/Spermatogenesis2018

## Shiny server

To visualize the different samples named in this study, we set up a shiny app which can be accessed via: https://marionilab.cruk.cam.ac.uk/SpermatoShiny

## Author Contributions

C. E., N.E., J.C.M., D.T.O. designed experiments; C.E. performed all experiments presented in the manuscript and interpreted the data; N.E. performed computational analysis and interpreted the data; C.P.M.J. performed preliminary experiments and provided technical assistance; C.E., N.E., J.C.M., D.T.O wrote the manuscript. All authors commented on and approved the manuscript.

## Acknowledgements

We thank the CRUK-CI Genomics (Paul Coupland and Katarzyna Kania), Flow Cytometry (Jelena Markovic-Djuric and Richard Grenfell) and Histopathology cores and the Biological Resources Unit for technical assistance. We thank Michael Morgan for technical help and Aaron Lun for providing help on the CUT&RUN analysis. We thank Steven Henikoff for kindly providing purified proteinA-MNase protein as well as yeast-spike DNA for CUT&RUN experiments. This research was supported by European Molecular Biology Laboratory (N.E., J.C.M.), Cancer Research UK (C.E., C.P.M.J., D.T.O., J.C.M.), the Wellcome Sanger Institute (C.P.M.J., J.C.M.), the Wellcome Trust (C.E., J.C.M. - grant 105031/Z/14/Z) and the European Research Council (D.T.O. - grant 615584).

